# Postharvest partial dehydration of blueberries enhanced blueberry wine aroma via upregulating phenylalanine metabolism and terpene biosynthesis

**DOI:** 10.1101/2024.01.29.577155

**Authors:** Yu Wang, Qi Zhang, Qin Yang, Chen Bian, Shu-Qin Huang, Lu-Lu Zhao, Ya-Qiong Huang, Shan-Shan Shen, Qi Chen, Hai-Wei Zhang, Xue-Ling Gao

## Abstract

Postharvest partial dehydration of blueberries can enhance blueberry wine aroma, while the underlying mechanisms remain unclear. In this study, the key odor-active volatiles in blueberry wines fermented from dehydrated blueberries (30% weight loss) were identified via aroma extract dilution analysis. Results showed that increased levels of phenylalanine-derived compounds such as phenylethanol, and terpenes such as linalool and geraniol, primarily led to the enhancement of sweet, floral and fruity aromas of blueberry wines. Postharvest partial dehydration increased the contents of these compounds, which could be linked to the upregulation of *VcGOT2* and *VcPAR* involved in phenylalanine metabolism, and the upregulation of *VcDXS*, *VcHDR* and *VcTPS* involved in terpene biosynthesis. Notably, the upregulated *VcTPS* encoded a monoterpene synthase responsible for producing linalool. These findings provided insight into the impact of postharvest dehydration on phenylalanine and terpene metabolism in blueberries, offering a reference for improving blueberry wine aroma through postharvest partial dehydration techniques.

## 1. Introduction

Blueberry is known for its great health potential and unique flavor, and its production has been rapidly increasing in recent years in China. However, blueberry is perishable and has a relatively short shelf life because of high water content and thin pericarp (Yuan et al., 2020). As blueberry production increases, blueberry processing has attracted rising attention, aiming at avoiding the massive damage and spoilage and improving the economic value and availability of blueberries. Blueberry wine remains abundant bioactive compounds and typical flavors of blueberries, satisfying consumers’ preference of fruit wines, in particular the health potential and diverse flavor. However, the weak pleasant aromas and less complex flavor of blueberry wines limit the development of blueberry winemaking (Wang et al., 2023a). Our previous research suggested that postharvest partial dehydration of blueberries substantially enhanced aroma of blueberry wines, especially sweet, floral and fruity odors, showing the possibility in innovating blueberry wine products and improving their flavor quality (Wang, et al., 2023b). However, the contribution of odor-active components to the aroma enhancement of blueberry wines should be further investigated.

Phenylalanine-derived volatiles in blueberry wines primarily include phenylacetaldehyde, phenylethanol, phenethyl acetate, etc., mainly contributing to honey/sweet and rose-like odors (Lan et al., 2019). Theoretically, phenylalanine can be converted to phenylacetaldehyde by aromatic amino acid decarboxylase (AADC) or phenylacetaldehyde synthase (PAAS), and further be transformed into phenylethanol by phenylacetaldehyde reductase. Phenylethanol can be transesterified to phenethyl acetate via the action of alcohol acyltransferase (AAT). Notably, albeit several key genes and enzymes involved in the formation of phenylalanine-derived volatiles have been characterized in some species (Maoz, Lewinsohn, & Gonda, 2022) such as tomato fruits, *Petunia* and rose flowers etc., the structural and molecular regulatory mechanisms leading to the phenylalanine-derived volatiles *in-planta* are still unknown in many cases such as blueberry. It has been reported that phenylacetaldehyde and phenylethanol have important biological functions in plants, and their synthesis can be induced by multiple stresses (Tieman et al., 2006). Our previous study found that phenylalanine content significantly decreased in partially dehydrated blueberries, while phenylacetaldehyde, phenylethanol and phenethyl acetate levels increased (Wang et al., 2023b). Similarly, Chen et al. (2019) reported that off-vine grapes underwent postharvest dehydration retained higher content of phenylethanol. Therefore, we proposed that postharvest dehydration promoted phenylalanine metabolism, favoring the biosynthesis of benzeneacetaldehyde and phenylethyl alcohol and phenethyl acetate in blueberries, consequently leading to the enhancement of sweet note of blueberry wine. However, the regulation mechanism of postharvest partial dehydration on phenylalanine metabolism remains unclear, in particularly the expression profiles of key genes involved.

The floral and fruity aromas of blueberry wines are highly associated with the terpenoids, which is largely determined by terpenes derived from blueberry fruits. Previous studies have identified the odor-active components of blueberries from different cultivars (Du et al., 2011; Du & Rouseff, 2014; Pico, Gerbrandt, & Castellarin, 2022). The primary odorant terpenes in blueberries included linalool, α-terpineol, geraniol, limonene, eucalyptol, and carveol, etc. Recently, Ferrão et al. (2022) combined information from consumer sensory perception of blueberry flavor and the knowledge of volatile compounds presented in blueberry fruits, revealing that the terpenes largely determined the aroma typicality and overall preference of blueberries and their derivative products. Liu et al. (2019) confirmed that terpenes derived from blueberries can be gradually extracted into the blueberry wines, shaping the terpenoid profile of blueberry wines. Our previous study also suggested that the enhancement of floral and fruity aroma of blueberry wines produced from the dehydrated blueberries was highly associated with the increased terpene content (Wang et al., 2023b).

The terpene increase in dehydrated berries and the resultant blueberry wines can not only be attributed to the concentration effects but also the upregulated terpene biosynthesis induced by water loss (Mencarelli & Bellincontro, 2020; Sanmartin et al., 2021). Terpenes in blueberries are produced from two biosynthetic pathways, namely the cytosol-localized mevalonic acid (MVA) and the plastid-localized 2-C-methyl-D-erythritol-4-phosphate (MEP) pathways. The key odorant monoterpenes are derived from MEP pathways catalyzed by a series of key enzymes, including 1-deoxy-D-xylulose-5-phosphate synthase (DXS), 1-deoxy-D-xylulose-5-phosphate reductoisomerase (DXR), 1-hydroxy-2-methyl-butenyl 4-diphosphate reductase (HDR), and terpene synthase (TPS). Terpenes are presented in free or glycosylated forms in blueberries, and the latter is formed by the action of glycosyl transferase (Lin, Massonnet, & Cantu, 2019). Several previous research has reported that postharvest dehydration upregulated the expressions of *TPS* genes in grapes (Zenoni et al., 2016; Zenoni et al., 2020), subsequently promoted the accumulation of terpenes. In blueberries, the biological function of key candidate structural genes involved in the terpene biosynthesis needs to be further reviewed, and the regulatory effects of postharvest dehydration on their expressions remains unclear.

In this study, we first evaluated the effects of postharvest partial dehydration on aroma profiles of blueberry wines, and further identified odor-active components by aroma extract dilution analysis. Subsequently, we tracked the dynamic changes of phenylalanine-derived volatiles and terpenes, and the responses of relevant genes to postharvest dehydration in blueberries, aiming to identify the key genes controlling the metabolite changes. And on this basis, we studied the potential biological function of a key terpene synthas. This study would provide new insights into phenylalanine metabolism and terpenoid metabolism in blueberries during postharvest dehydration, assisting blueberry wineries to better apply postharvest dehydration technology to improve blueberry wine sensory.

## 2. Material and methods

### 2.1 Postharvest dehydration of blueberries

The fully ripened fresh blueberries (the whole berry turned blue) of *Vaccinium corymbosum* L. ‘Lanmei 1’ were manually harvested in a commercial blueberry orchard in Huaining, Anhui, China (N30°20’, E116°28’) in 2023. Berry samples were precooled in an air-conditioned room at approximately 20 °C for 30 min to remove the field heat. Subsequently, the blueberries were divided into six perforated boxes (1000 g per box). Each box was used as a replicate. Three boxes of fresh blueberries with 0% wight loss were directly used for blueberry winemaking as control (CK), and the rest were stored at 20 °C and ∼70% relative humidity for postharvest partial dehydration. During postharvest dehydration period, the weight loss and the health status of blueberries was monitored daily, the healthy blueberries were sampled every two days. Sampled berries were immediately frozen in liquid nitrogen for further analysis. When the weight loss of blueberries reached ∼30% (WL30), the dehydrated blueberries were used for blueberry winemaking following the same procedure as CK.

### 2.2 Laboratory scale fermentation of blueberry wines

Firstly, the blueberries were manually crushed, and 50 mg/kg pectinase and 50 mg/L equivalent SO_2_ (as potassium metabisulfite) were subsequently added to the blueberry must. The blueberry must was put into a sterilized glass fermenter (500 mL) and maintained at room temperature in darkness for 24 hours, allowing for maceration and pectin hydrolysis. The total soluble solids of the blueberry must were adjusted to 20 °Brix by the addition of sucrose prior to alcoholic fermentation, followed by an inoculation with activated *Saccharomyces cerevisiae* Zymaflore RX60 (Laffort, Bordeaux, France). The fermentation temperature was maintained at 25 °C in an incubator. When the total sugar of blueberry wine was below 4 g/L, the pomace was separated, and the blueberry wine was filtered with addition of 30 mg/L equivalent SO_2_ (as potassium metabisulfite).

### 2.3 Determination of physiochemical compositions

A digital pocket handheld refractometer (Atago PAL-1, Tokyo, Japan) and a pH meter (Mettler-Toledo S220, Greifensee, Switzerland) were used to determine total soluble solids (TSS) and pH values, respectively. Sugar content, titratable acidity, ethanol level, and total SO_2_ in blueberry juice and wine were determined according to “Compendium of International Methods of Analysis of Wines and Musts (Edition 2020, Volume 1)” established by the International Organization of Vine and Wine. Total phenol content was determined by the Folin-Ciocalteu method and was expressed as gallic acid equivalents. Total anthocyanin content was determined by pH-differential method and was expressed as equivalent cyanidin-3-glucoside. The color of blueberry juice and wines was expressed by the CIELab parameters, including brightness (L*), red/green component (a*) and yellow/blue component (b*). A 10° standard observer and a standard light source D65 were used. Absorbance values of the wine samples were recorded at 450, 520, 570 and 630 nm by UV-vis spectrophotometer (Perkin Elmer Lambda35, Waltham, USA).

### 2.4 Aroma assessment of blueberry wines

A sensory panel consisted of ten assessors (eight females and two males) from the School of Tea and Food Science & Technology at Anhui Agricultural University, and they aged between 21 and 25 with an average of 23. Each panel member was trained for at least 40 hours for odor and intensity recognition using a “Le Nez du Vin” 54 Wine Aroma kit (supplied by Ease Sent Wine Education Co., Ltd, Beijing, China). The wine samples (15 mL per glass) were served in standard wine-tasting glasses labeled with a three-digit random code, and they were presented to the assessors in random order. Firstly, each assessor was asked to provide odor descriptors of blueberry wines as much as possible. Subsequently, a group discussion was performed to compile the descriptors with similar interpretations and remove the descriptors with less agreement. The final odor descriptors of blueberry wines included floral, fruity, honey, butter-like, toasty, herbaceous, jam-like, and hawthorn-like. Finally, the assessors were instructed to rate the intensity of each attribute on a continuous 0-5 scale (0, none; 1, very weak; 2, weak; 3, medium; 4, strong; 5 very strong) after sniffing blueberry wine samples. Each sample evaluation was performed three times.

### 2.5 Aroma extract dilution analysis of blueberry wines

Volatile compounds in blueberry wines were extracted using a headspace solid phase microextraction (HS-SPME) method. Samples were agitated at 2000 r/min for 5 min at 50 °C, and a 2 cm pre-conditioned (250 °C for 1 h) DVB/CAR/PDMS 50/30 μm SPME fiber (Supelco, Belletfonete, USA) was inserted into the headspace of the vial to absorb volatiles at 50 °C for 30 min. Afterward, the SPME fiber was inserted into the gas chromatography (GC) injector for 5 min at 230 °C (Wang et al., 2023c).

Blueberry wine samples were analyzed using an Agilent 7890B-5977B gas chromatograph/mass spectrometry (Agilent, Santa Clara, USA) coupled with an Gerstel ODP3 olfactometry system (Gerstel, GmbH&Co.KG, Germany). The volatile components were separated by a DB-5 capillary column (30 m × 0.25 mm × 0.25 μm, Agilent, CA, USA) using high-purity helium as carrier gas with a 1.5 mL/min flow rate. The flow of the carrier gas was split between a mass detector and the olfactometry system in a 1:1 ratio. The injection temperature was set at 230 °C. The oven temperature was programmed at the successive temperature: 40 °C for 5 min, increased to 180 °C at a rate of 3 °C /min, and then increased to 300 °C at a rate of 30 °C /min and held for 2 min. Mass spectra were generated in the electron ionization mode at 70 eV with a scan range of m/z 35-300. The ion source and quadrupole temperature were set at 230 °C and 150 °C, respectively.

The extracts were stepwise diluted by setting the split ratio of the oven at 1:1 ratio up to a 1024 dilution factor until the odor is not felt at the sniffing port. Each diluted sample was repeatedly sniffed by three sniffing panel members until no odorous compounds were detected in the olfactometry system. The flavor dilution factor (FD) for each volatile compound in blueberry wines was defined as the final dilution at which the odor-active zone could be smelled by two or more panelists.

### 2.6 Determination of volatile compounds in blueberries and blueberry wines

Since volatile compounds in blueberries were presented in free and bound forms. In this study, the total volatile compounds (free + bound) were analyzed followed by a direct enzyme hydrolysis. Briefly, one g of fine grinded blueberry sample was diluted with 9 mL of citrate buffer (0.2 M, pH 5.0) in a 20 mL vial. Subsequently, 100 μL Rapidase AR2000 (DMS Food Specialties, Delft, Netherlands) enzyme solution (0.07 g/mL) were added into the mixture, which was incubated at 40 °C for 16 h in a tightly capped vial. After the solution was cooled to room temperature, 3 g NaCl and 20 μL internal standard (22.71 μg/mL 3-nonanone) were added into the mixture. The extraction of volatile compounds from blueberry and blueberry wines were performed following the same HS-SPME method as described above.

Volatile compounds were analyzed using an Agilent 6890 GC coupled with an Agilent 5975C MS (Agilent, Santa Clara, USA). Volatile compounds were separated by an HP-5MS capillary column (30 m × 0.25 mm, 0.25 μm thickness, J&W Scientific, Folsom, USA). The flow rate of high-purity helium carrier gas was 1.0 mL/min. The injection temperature was set at 230 °C in splitless mode. The oven temperature was programmed at the successive temperature: 35 °C for 5 min, increased to 130 °C at a rate of 4 °C/min and held for 3 min, and then increased to 230 °C and held for 1 min. The ion source and quadrupole temperature were set at 230 °C and 150 °C, respectively. The ionization voltage was 70 eV. Full scan mode was applied to collect electron ionization mass data from m/z 30-350. The identification and quantification of volatile compounds in blueberries and blueberry wines were followed the methods described by Wang et al (2023b).

### 2.7 Extraction of RNA and RT-qPCR analysis

Total RNA of blueberries was extracted using a Spin Column Plant Total RNA Purification Kit (Sangon Biotech, Shanghai, China). The concentration and purity of RNA was determined using a Thermo NanoDrop ND-2000 spectrophotometer (Wilmington, DE, USA). The integrity of RNA was verified via electrophoresis on 1.0 % agarose gel. cDNA library was obtained via reverse transcription using the MightyScript First Strand cDNA Synthesis Master Mix Kit (Sangon Biotech, Shanghai, China).

Quantitative real-time PCR was conducted to assess the relative expressions of genes involved in phenylalanine metabolism and terpene biosynthesis with a Bio-Rad CFX96 instrument (Bio-Rad, Shanghai, China). The gene-specific primers used for qRT-PCR was listed in Table S1, and *VcUBC28* was applied as the reference gene (Die & Rowland, 2013). Each qRT-PCR reaction (20 μL) contains 1 μL of cDNA template, 0.4 μL of 10 mM forward primer, 0.4 μL of 10 mM reverse primer, 8.2 μL of ddH_2_O, and 10 μL of 2 × SGExcel FastSYBR qPCR Mix solution (Sangon Biotech, Shanghai, China). The cycling conditions were as follows: 95 °C for 3 min, followed by 40 cycles of 95 °C for 5 s, 60 °C for 20 s. Relative fold differences were calculated using the 2^−ΔΔCt^ method.

### 2.8 Identification and classification of *VcTPS* genes in the *Vaccinium corymbosum*

PF01397 and PF03936, representing the TPS N-terminal domain and the TPS C-terminal domain from PFAM1, respectively, were used as queries to search the recent *Vaccinium corymbosum* L. protein database (Colle et al., 2019). An HMM (Hidden Markov Model) search was used in this study with an e-value cut at 10^−3^ by using Tbtools softwar. To avoid missing potential TPS genes, 32 known TPS sequences from *Arabidopsis thaliana* were also used to screen the *Vaccinium corymbosum* protein database using BLASTP (built-in Tbtools software). The candidate TPS genes were checked manually by NCBI batch CDD search and InterPro online tools to verify putative full-length TPS genes, and TPS genes with incomplete/partial conserved domain or lacking either PF03936 or PF01397 were excluded. The molecular weight (Mw), isoelectric points (pI), aliphatic index (AI), grand average of hydrophobicity (GRAVY), and instability index (II) of the TPS proteins were predicted by the ExPASy database (Artimo et al., 2012).

To classify the evolutionary relationships of the *VcTPS* gene family, the TPS protein sequences of *Arabidopsis thaliana*, tomato (*Solanum lycopersicum*), grape (*Vitis vinifera*), and blueberry (*Vaccinium corymbosum*) were aligned using the CluastW algorithm and MEGA11 software. Phylogenetic analysis based on amino acid sequence alignment was performed using the neighbor-joining (NJ) method with Bootstrap tests on 1000 resamples, and maximum likelihood (ML) was considered for more reliable phylogenetic analysis. The final phylogenetic relationship was based on the results obtained from the two methods and visualized by iTOL online software.

### 2.9 Protein expression in *E. coli* and terpene synthase assays

The ORF of *VcTPS15* was synthesized and ligated to into pET-22b vector. Subsequently, the recombinant plasmid was transformed into *Escherichia coli DH5*α competent cells (Abiocenter, Wuxi, China). Positive colonies were selected and fully sequenced to assess identity by DNA amplification. Selected clones were transformed to *E. coli* BL21 (DE3) strain (Abiocenter, Wuxi, China). For induction of protein expression, single colonies were inoculated in 5 mL LB medium (Abiocenter, Wuxi, China) and grew overnight at 37 °C. Aliquots of 200 μL were inoculated in 200 mL fresh LB medium, and cultures were grown at 37 °C with shaking at 250 rpm until OD_600_ = 0.6. For induction of recombinant protein expression, IPTG (isopropyl β-D-thiogalactopyranoside) was first added to a final concentration of 0.5 mM and cultures were maintained at 18 °C for 16-18 h with shaking. The cells were harvested by centrifugation at 1,2000 rpm for 15 min at 4 °C. The precipitate was resuspended in lysis buffer (50 mM Tris, 0.5 M NaCl) and disrupted by ultrasonic treatment (work 3 s, off 2 s) for 15 min. Cell debris were removed by centrifugation at 1,2000 rpm for 30 min at 4 °C. The precipitate was resolved into assay butter (50 mM Tris, 0.15 M NaCl, and 8 M urea, pH 7.4), and protein was purified by immobilized affinity metal chromatography and then examined by SDS-PAGE. Enzyme assays were performed in 1 mL assay buffer (30 mM HEPES, 5 mM DTT, 25 mM MgCl_2_, pH 7.5) containing 60 μM geranyl diphosphate (GPP), and 10 μg purified VcTPS15 protein. The mixture was incubated at 30 °C for one hour and then 45 for 15 min (Falara et al., 2011). Subsequently, the reaction products were determined by SPME-GC/MS following the procedure described above.

### 2.10 Statistical analysis

One-way analysis of variance (ANOVA) employing Duncan’s multiple range test at p < 0.05 was performed using “agricolae” in the R environment (4.0.1). Sensory assessment data was processed using PanelCheck software v1.4.2 (Nofima, Norway). Barplots and line plots were conducted using Origin 2021 (Originlab, USA). Heatmap was prepared using the “ComplexHeatmap” package in the R environment (4.0.1).

## 3. Results and discussions

### 3.1 Physicochemical parameters of blueberries and blueberry wines

Table S2 showed the effects of postharvest partial dehydration on the basic physiochemical parameters of ‘Lanmei 1’ blueberries and resulting wines. As expected, total soluble solids and sugar content of blueberries were significantly increased by postharvest dehydration treatment due to the concentration effects. In agreement with our previous findings, the titratable acidity showed a marked decrease in juice and final wines produced from dehydrated blueberries. These decreases could be attributed to the enhanced conversion and catabolism of malic acid and citric acid, two primary organic acid presented in blueberries, induced by cellular oxidative decomposition or gluconeogenesis. Notably, total anthocyanins and phenols in dehydrated blueberries and resulting wines were significantly higher than controls. Beside concentration effects, postharvest dehydration might also induce the upregulation of key genes involved in anthocyanin biosynthesis. Additionally, dehydrated berries might lead to the internal degradation of the pericarp cell layer, consequently enhancing the extractability of anthocyanins and phenols. However, when total anthocyanin content was expressed on a per berry basis, it was decreased in WL30 blueberries. Besides, when total phenol content was expressed on a per berry basis, no significant differences were observed between WL30 blueberries and controls. These results indicated that increases in total anthocyanins and phenols in blueberry wines fermented from dehydrated blueberries could be mainly ascribed to the concentration effects caused by water loss. The color of blueberry wine was largely dependent on the phenol components, especially anthocyanins. A significant shift towards redness and darkness was observed in blueberry wines produced from dehydrated berries. There were no significant differences in residual sugar, ethanol level and SO_2_ content between WL30 blueberry wines and controls.

### 3.2 Aroma profiles of blueberry wines

Eight descriptors were given by the trained panelist to shape the blueberry wine aroma profile, namely floral, fruity, honey, butter-like, toasty, herbaceous, jam-like, and hawthorn-like (Figure 1). Postharvest partial dehydration of blueberries significantly increased the intensity of floral, fruity, honey and jam-like aromas but decreased herbaceous aroma intensity (*p* < 0.01). In agreement with our results, previous research also reported that wines derived from the dehydrated grapes were characterized by stronger honey/sweet, floral, and fruity notes (Lan et al., 2019; Ma et al, 2021). The odor-active compounds, contributing to these typical aromas needed to be identified, and the regulatory effects of postharvest dehydration on these compounds in blueberries should be further investigated.

**Figure 1.**
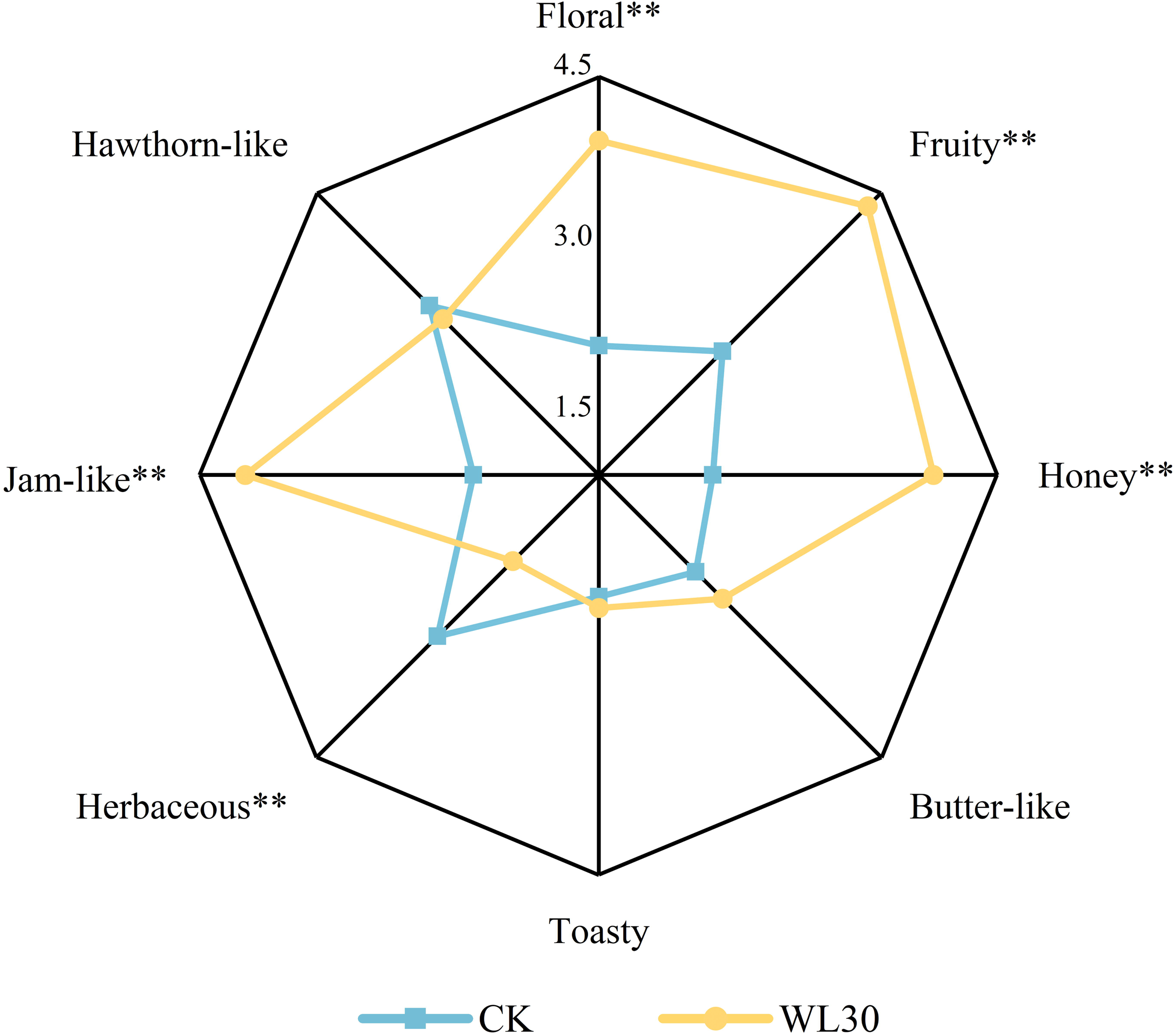
Sensory analysis of blueberry wines fermented from fresh (CK) and partially dehydrated blueberries (WL30). Significance levels were analyzed by LSD as follows: * *p* < 0.05, ** *p* < 0.01.

### 3.3 Identification of key odor-active volatile compounds in blueberry wines

A total of 37 odor active regions were detected in WL30 and control blueberry wines by the olfactory senses of two or more experienced judges (Table 1). A total of 29 odorants were further identified by comparing the retention indices (RIs), mass spectrometry, and odor quality with the corresponding reference compounds. These compounds can be divided into 7 groups, including esters, terpenes, phenylalanine derivatives, and volatile phenols. WL30 and CK blueberry wines had similar odorant compositions, sharing 32 common odor active regions. Notably, 1,4-cineol, *p*-tolualdehyde, *cis*-β-farnesene, and ethyl isobutyrate were only detected in WL30 but not in controls (Table 1), mainly contributing to floral, fruity and sweet aromas.

**Table 1.** Key odor-active compounds identified in blueberry wines fermented from fresh blueberries and partial dehydrated blueberries by AEDA-GC/MS-O.

FD factors were calculated to evaluate the contribution of each odor-active compounds to blueberry wine aroma. In CK, only 2 odor substances, linalool and γ-terpinene, had FD values equal to 1024, lower than those in WL30 blueberry wines. By contrast, there are 17 odor substances in WL30 with FD values greater than or equal to 1024 (Table 1), including linalool (>1024; floral), β-pinene (>1024; floral, sweet), γ-terpinene (>1024; floral, sweet), β-ocimene (>1024; floral), *cis*-geraniol (>1024; floral, mint-like), phenylethanol (>1024; rose), phenyl acetate (>1024; honey-like, sweet), *trans*-β-damascenone (1024; floral, honey-like), camphor (1024; floral, mint-like), *endo*-borneol (1024; floral, mint-like), 4-ethylguaiacol (1024; jam-like, sweet), *p*-menth-1-en-9-ol (1024; mint-like), engenol (1024; lilac), isoamyl acetate (1024; banana-like, sweet), ethyl hexanoate (1024; fruity), and ethyl decanoate (1024; fruity). FD factors of all these compounds in WL30 blueberries were increased by at least 100% compared to CK (Table 1). Additionally, most of these compounds contribute to three aroma attributes of blueberry wines, namely sweet, floral, and fruity notes.

It was noted that phenylalanine derivatives such as phenylethanol and phenyl acetate, and terpenes such as linalool, β-pinene, γ-terpinene, β-ocimene and *cis*-geraniol had the highest FD factors, higher than 1024, in WL30 blueberry wines (Table 1). In general, these results revealed the large differences in FD factors between blueberry wines produced from dehydrated and fresh blueberries, which were primarily caused by terpenes and phenylalanine derivatives, contributing to the sweet, floral, and fruity aromas. Consistently, several studies have also reported that the honey/sweet, tropical fruit and floral aromas of wines produced from dehydrated grapes were characterized by phenylacetaldehyde, phenethyl alcohol, phenyl acetate, β-damascenone, linalool and geraniol (Lan, et al., 2019; Ma, Xu, & Tang, 2021; Moreno et al., 2008; Urcan et al., 2017). Therefore, we mainly determined phenylalanine-derived compounds and terpenes in blueberries and resulting wines in the subsequent analysis.

### 3.4 Relative quantitation analysis of phenylalanine-derived compounds and terpenes in blueberry wines

Seven phenylalanine-derived compounds were identified in blueberry wines via SPME-GC/MS analysis, and the concentrations of all these compounds were significantly increased by postharvest dehydration (Figure 2A). Among these compounds, phenethyl alcohol, phenethyl acetate and methyl salicylate were also detected via GC/MS-O, showing higher FD factors in WL30 blueberry wines than in controls (Table 1). The rOAVs of methyl salicylate, phenylethanol, phenethyl acetate, ethyl phenylacetate and phenylacetaldehyde was increased by at least 47% in WL30 blueberry wines, greater than 0.1, especially that the concentrations of ethyl phenylacetate and phenylacetaldehyde exceeded their corresponding olfactory thresholds in WL30 (Table S3). Most of these compounds imparted rose-like and honey/sweet aromas, thus the above findings confirmed the positive roles of postharvest dehydration in enhancement of floral and sweet aromas of blueberry wines via increasing phenylalanine derivative content. Consistently, several previous studies also characterized phenylalanine derivatives as key active odorants in wines produced from dehydrated berries (Ossola, et al., 2017). The formation of phenylalanine derivatives in wines could be originated either from those precursors presented in fruits, or from the degradation of phenylalanine during fermentation via yeast metabolism. The associations between the effects of postharvest dehydration on phenylalanine metabolism in blueberries and the final phenylalanine derivatives in blueberry wines needed to be further investigated.

**Figure 2.**
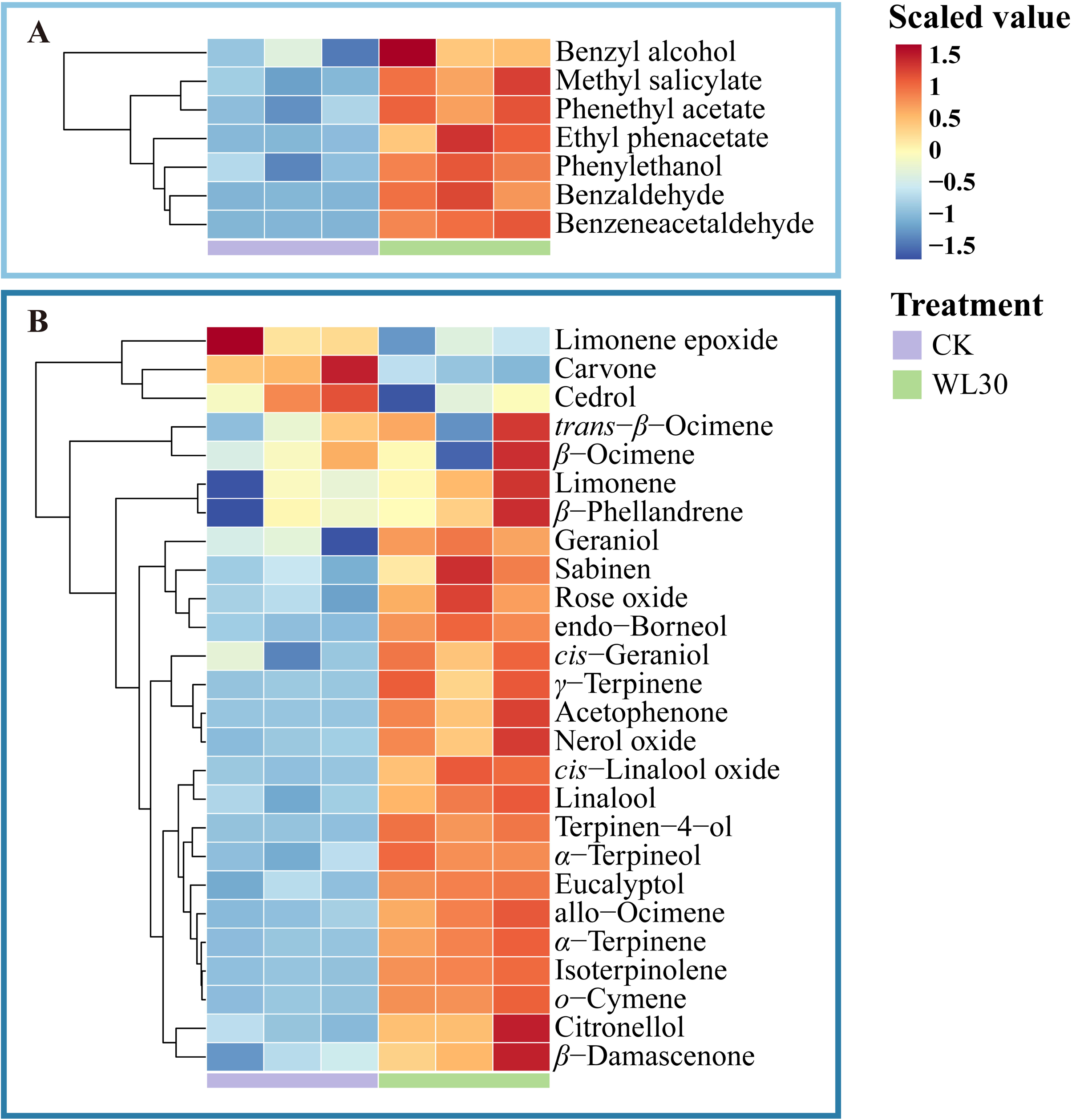
Heatmap visualization of the concentrations of phenylalanine-derived volatiles (A) and terpenes (B) in blueberry wines fermented from fresh (CK) and partially dehydrated blueberries (WL30).

A total of 26 terpene compounds were identified in blueberry wines. It was noted that the concentrations of 19 terpenes in WL30 were significantly higher than in controls (Figure 2), showing the marked positive effects of postharvest partial dehydration on terpenes in blueberry wines. Linalool was the most abundant terpenes, followed by α-terpineol, carvone, citronellol, and geraniol (Table S3). Among these compounds, only α-terpineol and carvone were not detected by GC/MS-O, which might be linked to their olfactory thresholds. According to rOAVs, linalool exceeded it olfactory thresholds in both CK and WL30 blueberry wines, and its rOAV was increased by 5.5 by postharvest dehydration. The rOAV of geraniol in WL30 were higher than 1 but not in CK, and rOAV of citronellol in WL30 was increased by 1.2 folds, reaching to 0.91. The highest rOAV was observed on β-damascenone because of its extremely low olfactory thresholds, and its rOAV was increased by 51% by postharvest dehydration (Table S3). Consistently, several studies also reported that sweet wines and ice wines produced from dehydrated grapes possessed higher content of β-damascenone than those made from fresh grapes harvested at maturity (Bowen & Reynolds, 2012; Lan, et al., 2019; Qian et al., 2024). The most significant change was recorded for terpinen-4-ol which was increased by 63 folds in WL30 blueberry wines (Table S3). In agreement with our findings, previous studies have characterized terpinen-4-ol as a marker of grape postharvest dehydration (Negri et al., 2017; Shmuleviz et al., 2023). Furthermore, the rOAV of terpinen-4-ol in CK was lower than 0.1 but higher than 1 in WL30 blueberry wines, suggesting that the aroma contribution of terpinen-4-ol was substantially enhanced in blueberry wines produced from dehydrated berries (Table S3). In general, the above findings revealed that linalool, geraniol, citronellol, β-damascenone, and terpinen-4-ol were the key odor-active terpenes in blueberry wines, which was consistent with previous studies (Qian et al., 2021; Sater, Bizzio, Tieman, & Muñoz, 2020). Higher rOAVs of these compounds in WL30 blueberry wines were in accordance with the increases in corresponding FD factors via GC/MS-O analysis, contributing to the enhancement of floral and fruity notes of blueberry wines produced from dehydrated berries.

### 3.5 Dynamic changes phenylalanine-derived compounds in blueberries during postharvest dehydration process

Four phenylalanine-derived compounds were detected in blueberries, including benzyl alcohol, benzaldehyde, phenylethanol, and ethyl phenylacetate (Figure 3, Table S4), which were also found in the resulting blueberry wines (Figure 2). The content of benzyl alcohol and benzaldehyde showed similar increasing trends during postharvest dehydration process either on a per fresh weight basis or on a per berry basis (Figure 3, Table S5). The highest concentrations of benzyl alcohol and benzaldehyde were observed in WL30 blueberries, increased by 3.6 and 29.7 folds compared to fresh blueberries on a per fresh weight basis, respectively (Table S4). These results indicated that the increases in benzyl alcohol and benzaldehyde can not only be attributed to concentration effects but also upregulated biosynthesis. Benzyl alcohol and benzaldehyde could be derived from the β-oxidation of cinnamoyl-CoA, the Co-A ligation product of cinnamic acid which was converted from phenylalanine catalyzed by phenylalanine ammonia-lyase (PAL). However, in blueberry wines, the rOAVs of benzyl alcohol was lower than 0.1. Besides, neither benzyl alcohol nor benzaldehyde was detected via GC/MS-O analysis, showing their limited contributions to blueberry wine aroma.

**Figure 3.**
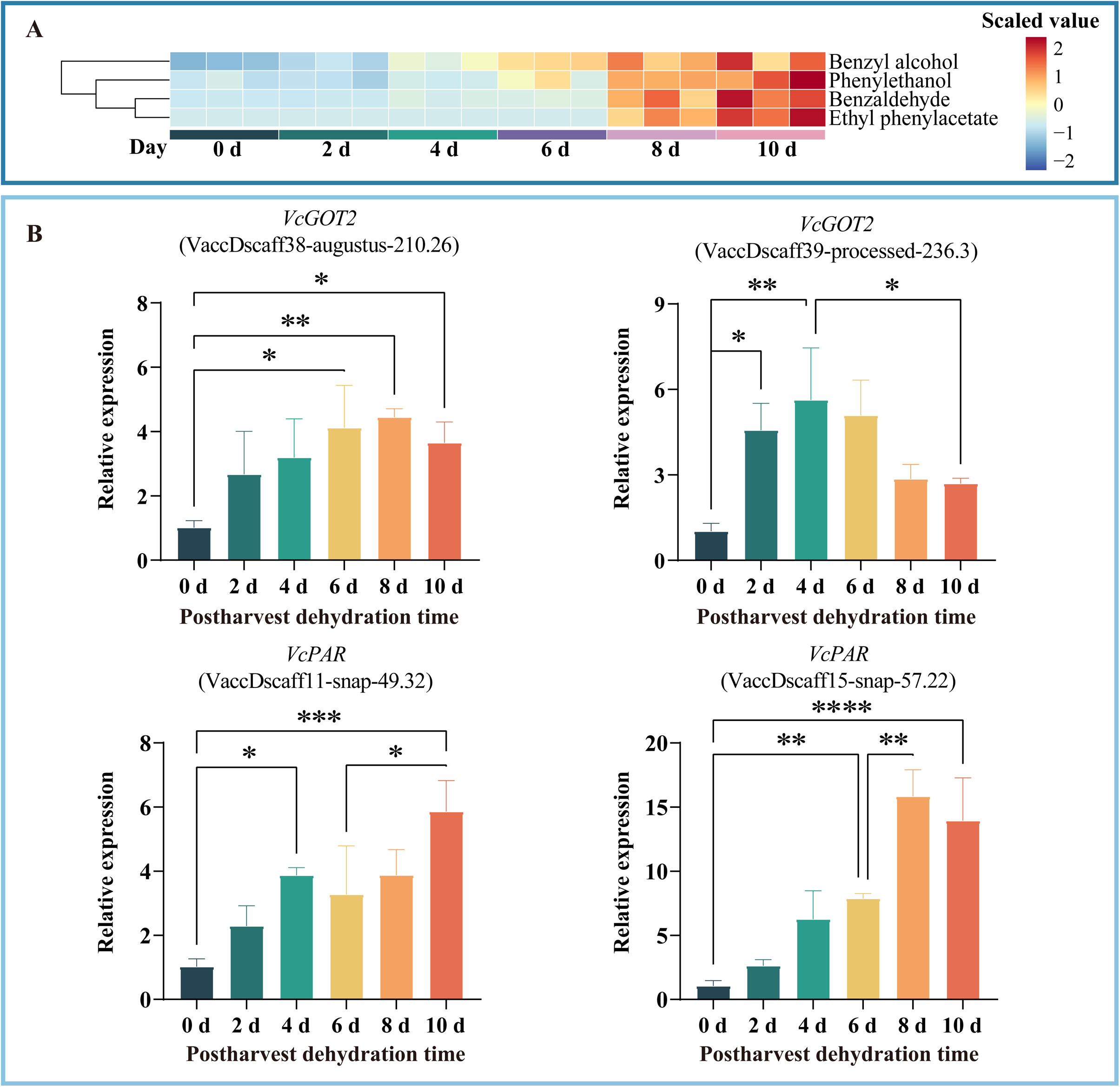
Heatmap visualization of the dynamic changes in the concentrations of phenylalanine-derived volatiles (A), and the dynamic changes in the relative expressions of *VcGOT2* and *VcPAR* genes involved in phenylalanine metabolism in blueberries during postharvest dehydration (B). Significance levels were analyzed by LSD as follows: * *p* < 0.05, ** *p* < 0.01, *** *p* < 0.001, **** *p* < 0.0001.

Phenylethanol has been characterized as a key odorant in blueberry wines by GC/MS-O and GC/MS analysis (Table 1, Table S3), and its higher content in WL30 blueberry wines could lead to the enhancement of sweet and floral notes. Phenylethanol in blueberry wines could be produced either from blueberries or from phenylalanine degradation via yeast metabolism during fermentation. Our previous studies suggested that phenylalanine content in dehydrated blueberries was significantly decreased by postharvest dehydration (Wang et al., 2023b). Consistently, a previous study also reported a declining trend of phenylalanine content in ‘Beibinghong’ grapes during postharvest on-vine dehydration (Li et al., 2023). Therefore, we speculated that the higher phenylethanol content in WL30 blueberry wines could be primarily attributed to the increases in phenethyl alcohol in dehydrated blueberries. In this study, the content of phenylethanol was significantly increased at 6, 8, and 10 days of dehydration, increased by 3 folds in WL30 blueberries than in controls on a per fresh weight basis (Table S4). Similar results were also observed when phenylethanol content was expressed on a per berry basis (Table S5), indicating the upregulated phenylethanol biosynthesis.

Phenylethanol in plants is derived from the reduction of phenylacetaldehyde catalyzed by phenylacetaldehyde reductase (PAR). Phenylacetaldehyde could be produced via several pathways. One is from phenylalanine by phenylacetaldehyde synthase (PAAS) or aromatic amino acid decarboxylase, which has been functionally characterized in rose and *Petunia* flowers and *Arabidopsis* (Maoz, Lewinsohn, & Gonda, 2022). Another phenylacetaldehyde biosynthesis pathway is from phenylalanine via phenylethylamine catalyzed by aromatic amino acid decarboxylase and monoamine oxidase (MAO), which has been characterized in tomato fruits (Tieman et al., 2006). In addition, phenylethylamine can be converted to phenylpyruvate by aspartate aminotransferase (GOT2), and the latter can be transformed to phenylacetaldehyde by phenylpyruvate decarboxylase (PDC). Several studies also reported that phenylpyruvate were the precursors of benzaldehyde and benzyl alcohol (Wang, et al., 2019). In blueberries, the specific route of phenylacetaldehyde and phenylethanol biosynthesis remains unclear.

To investigate the further reason behind the increases in phenylethanol content in dehydrated blueberries, we determined the expression profiles of *VcGOT2* and *VcPAR* genes via qRT-PCR, since these genes have been characterized as key differentially expressed genes via transcriptome analysis between dehydrated and control blueberries (RNA-seq data has been deposited into NCBI with accession number PRJNA974910). Two *VcGOT2* genes, VaccDscaff38-augustus-210.26 and VaccDscaff39-processed-236.3, showed different expression trends during postharvest dehydration. The first one was significantly upregulated at 6, 8 and 10 days compared to fresh blueberries, while the expressions of the latter one drastically upregulated at 2 and 4 days of dehydration, significantly higher than controls, followed by a decreasing trend. The upregulation of *VcGOT2* genes during postharvest dehydration could favor the accumulation of phenylpyruvate, as a precursor of phenylacetaldehyde. The expressions of two *VcPAR* genes, VaccDscaff38-augustus-210.26 and VaccDscaff39-processed-236.3, gradually increased during postharvest dehydration. The first one was significantly upregulated from 4 to 10 days of dehydration, and its expression level was increased by over two folds at 10 d. The latter one was significantly upregulated from 6 to 10 days of dehydration, and its expression levels was increased by over five folds at 8 and 10 days of dehydration. The upregulated *VcPAR* genes in partially dehydrated blueberries would promote the conversion of phenylacetaldehyde to phenylethanol. Consistently, a previous study also reported that the postharvest spreading treatment upregulated the expressions of *CsPAR* genes in green tea, thus improving the production and release of phenylethanol (Yu et al, 2021). Overall, the above findings suggested that postharvest dehydration upregulated the expressions of *VcGOT2* and *VcPAR* genes, leading to the significant increases in phenylethanol level in blueberries, contributing to the enhancement of sweet and floral aroma of resulting blueberry wines.

### 3.6 Dynamic changes of terpenes in blueberries during postharvest dehydration process

A total of 20 terpenes were detected in blueberries (Table S4), and 12 of them were consistently identified in the final blueberry wines (Table S3). Among these compounds, linalool, geraniol, β-pinene, *o*-cymene, *trans*-β-ocimene, β-ocimene, *endo*-borneol and terpinene-4-ol were identified as the key odorants in blueberry wines via GC/MS-O and GC/MS analysis, mainly contributing to floral and fruity aromas. A gradual increasing trend was observed in the concentrations of these compounds, except for terpinene-4-ol, during postharvest dehydration on a per fresh weight basis (Figure 4, Table S4). These findings suggested that the higher terpene content in WL30 blueberry wines could be mainly attributed to the significant increases in corresponding terpenes in blueberries caused by postharvest partial dehydration.

**Figure 4.**
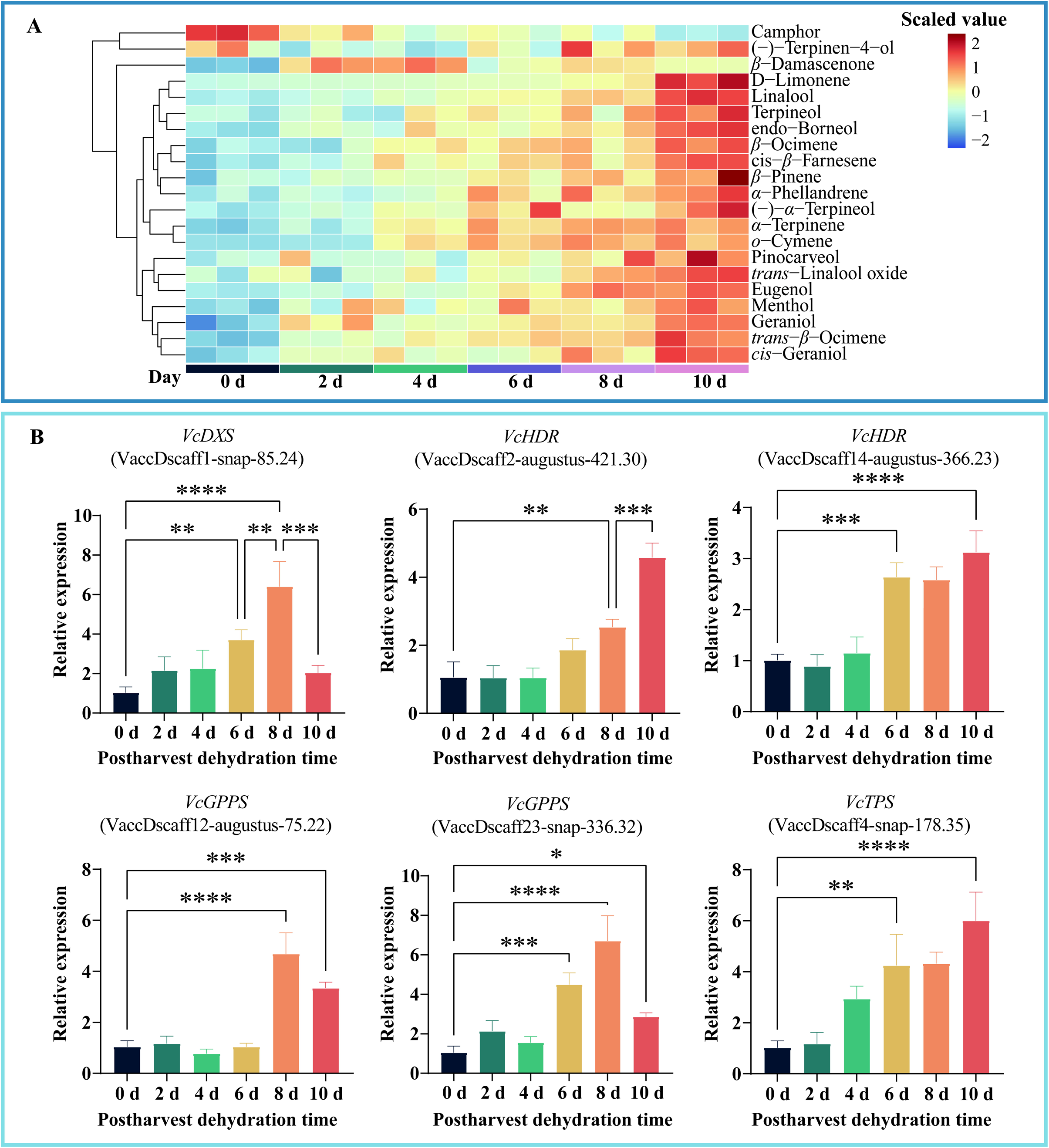
Heatmap visualization of the dynamic changes in the concentrations of terpenes (A), and the dynamic changes in the relative expressions of *VcDXS*, *VcGGPS*, *VcHDR*, and *VcTPS* genes involved in phenylalanine metabolism (B) in blueberries during postharvest dehydration. Significance levels were analyzed by LSD as follows: * *p* < 0.05, ** *p* < 0.01, *** *p* < 0.001, **** *p* < 0.0001.

Previous studies reported that the increases in terpene content was an integrative consequence of balance between concentration, biosynthesis, degradation, and oxidation, where concentration effect was more significant (Lan et al., 2016). The contents of linalool, geraniol, β-pinene, *o*-cymene and *trans*-β-ocimene expressed on a per berry basis in WL30 blueberries were also significantly higher than in controls (Table S5), showing that the increases of these terpenes in dehydrated blueberries can also be attributed to the upregulated biosynthesis besides concentration effects. Our previous study also confirmed that postharvest dehydration upregulated expressions of key genes involved in the MEP pathway in blueberries via transcriptome analysis, including *VcDXS*, *VcHDR*, *VcGGPS* and *VcTPS* (Wang et al., 2023). In this study, we conducted qPCR analysis to determine the dynamic changes of relative expressions of these genes during postharvest dehydration process. *VcDXS* and *VcHDR* are the key genes in the upstream of MEP pathway, controlling the biosynthesis of IPP and DMAPP, the precursors of monoterpenes and diterpenes/carotenoids respectively. The expressions of *VcDXS* (VaccDscaff-snap-85.24) were significantly upregulated at 6 and 8 days compared to fresh blueberries, followed by a significant downregulation. A significant upregulation of *VcHDR1* (VaccDscaff2-augustus-421.30) was observed at 8 and 10 days, and upregulation of *VcHDR2* (VaccDscaff14-augustus-366.23) was observed at day 6, 8, and 10 (Figure 4). *VcGGPS* is responsible for the biosynthesis of GGPP, the precursor of carotenoids, which can be further converted to norisoprenoids such as β-damascenone. The expressions of two *VcGGPS* genes were assessed, and *VcGGPS1* (VaccDscaff12-snap-75.22) was significantly upregulated at 8 and 10 days, while *VcGGPS2* (VaccDscaff23-snap-336.32) were upregulated at 6 and 8 days followed by a downregulation (Figure 4). TPS catalyze the final step of terpene biosynthesis, and it is regarded as a key rate-limiting enzyme. Our results showed that a *VcTPS* gene (VaccDscaff4-snap-178.35) was significantly upregulated after 6 days of dehydration (Figure 4). Notably, the above findings suggested that *VcDXS*, *VcHDR*, *VcGGPS* and *VcTPS* were upregulated on the mid/late dehydration stages (6 days later). The downregulation of *VcDXS*, *VcGGPS1*, and *VcGGPS2* at day 10 highlighted the importance of partial dehydration level, implying that terpene biosynthesis would cease in over-dehydrated berries where degradation and oxidation might be more significant.

Overall, the above findings suggested that postharvest partial dehydration could upregulate terpene biosynthesis in blueberries, consequently leading to the higher terpene levels in dehydrated blueberries and the resulting blueberry wines. The upregulation of *VcDXS* and *VcHDR* genes in the upstream of MEP pathway could partially explain the terpene increases, and the upregulation of *VcGGPS* genes could be linked to the higher β-damascenone content. Although we found a *VcTPS* gene which was significantly upregulated by postharvest dehydration, its final product remained unknown. The direct relationships between the upregulated *VcTPS* and the increased individual terpenes in dehydrated blueberries needed to be further investigated via functional characterization.

### 3.7 Identification and functional characterization of upregulated *VcTPS* gene in dehydrated blueberries

To investigate the function of upregulated *VcTPS* gene in dehydrated blueberries, we first conducted a bioinformatic analysis to identify and classify the *VcTPS* genes in blueberry. A total of 141 *VcTPS* genes were finally identified, and they were renamed according to their position on the chromosome as shown in Table S6. Among these genes, the upregulated *VcTPS* gene (VaccDscaff4-snap-178.35) in dehydrated blueberries in this study was defined as *VcTPS15*, with a 3640 bp in length, encoding 577 amino acids with 65.95 kDa molecular weight. The pI, aliphatic coefficients, instability index and hydrophilic coefficient of *VcTPS15* protein were 5.91, 80.97, 43.44 and −0.391, respectively, indicating that *VcTPS15* was an acidic, unstable protein, and was rich in aliphatic amino acids (Table S6).

Phylogenetic tree was constructed based on a multiple sequence alignment of the blueberry TPSs, *Arabidopsis thaliana* TPSs, *Solanum lycopersicum* TPSs, and *Vitis vinifera* TPSs, to discover the evolutionary relationships among the blueberry *VcTPS* genes. *VcTPS* genes in blueberry could be divided into five sub-families, and TPS-a, TPS-b, TPS-c, TPS-e/f and TPS-g sub-families contained 33, 49, 29, 12, and 18 *VcTPS* genes, respectively (Figure S1, Table S6). However, no TPS-d and TPS-h subfamily gene were identified in *Vaccinium corymbosum*. Consistently, previous studies reported that the TPS-d subfamily mainly distributed in gymnosperms species, and the TPS-h subfamily only existed in *Selaginella moellendorffii* (Chen, et al., 2011; Jiang et al., 2019). It has been reported that TPS-a genes mainly encode sesquiterpene synthases and diterpene synthases in various plants (Keilwagen et al., 2017), which was consistent with the function prediction of TPS-a proteins in this study (Table S6). It was noticed that TPS-b was the most expanded category in *Vaccinium corymbosum*, in parallel with the patterns in *D. officinale*, *V. planifolia*, *D. catenatum*, and *C. faberi*, having more genes in TPS-b (Yu et al., 2020; Huang et al., 2021). Furthermore, the angiosperm-specific TPS-b and TPS-g family encode monoterpene synthases, and the TPS-b and TPS-g members accounted for ∼50% of *VcTPS* genes in blueberry, which could be related to the more biosynthesis of monoterpenes and emission of floral scent in blueberry. In this study, the upregulated *VcTPS15* in dehydrated blueberries was classified as TPS-g sub-family (Figure S1, Table S6). We aligned the multiple sequence to analyze the conserved motifs of VcTPS15. The alignment showed that VcTPS15 contained R(X8)W, DDXXD and NSE/DTE motifs (Figure S2). In particularly, DDXXD and NSE/DTE motifs play an important role in the metal-dependent ionization of the prenyl diphosphate, and R(X8)W is essential in the cyclization of monoterpene synthase (Jiang et al., 2019). According to the protein function prediction, VcTPS15 was annotated as a monoterpene synthase.

An RNA-seq database of root, leaves (day and light), buds, flowers (FL and PF), and the fruits of different developmental stages (green fruit, pink fruit, and ripe ripe) was established to study the expression patterns of *VcTPS* genes in different organs (Figure S3). Notably, expressions of *VcTPS* gene in blueberry fruits showed developmental stage dependent and organ dependent patterns. Among these genes, *VcTPS15* was mainly expressed in pink and ripe fruit, and the highest expression was observed in ripe blueberries (Figure S3), suggesting that *VcTPS15* still played important roles in terpene biosynthesis in ripe blueberries. Our results further confirmed that *VcTPS15* would response to postharvest dehydration.

In order to investigate the biochemical function of terpene synthase, VcTPS15 protein was expressed in *E. coli*, and the terpene synthase activity of recombinant protein was determined. Figure 5 showed that VcTPS15 protein catalyzed GPP to β-linalool as the major product along with three minor monocyclic monoterpene products, including β-pinene, D-limonene and α-terpineol. The results indicated the biochemical function of VcTPS15 as monoterpene synthase. Consistently, the linalool, D-limonene and α-terpineol concentrations were significantly increased by postharvest partial dehydration, and the carry-on effects were also observed in the resulting blueberry wines. In particular, linalool was identified as a major aroma active compounds in blueberry wines contributing to floral aroma. Taken together, postharvest partial dehydration of blueberries upregulated the expressions of *VcTPS15*, leading to higher linalool concentrations in dehydrated blueberries and the resulting wines, thus enhancing the floral aroma of blueberry wines.

**Figure 5.**
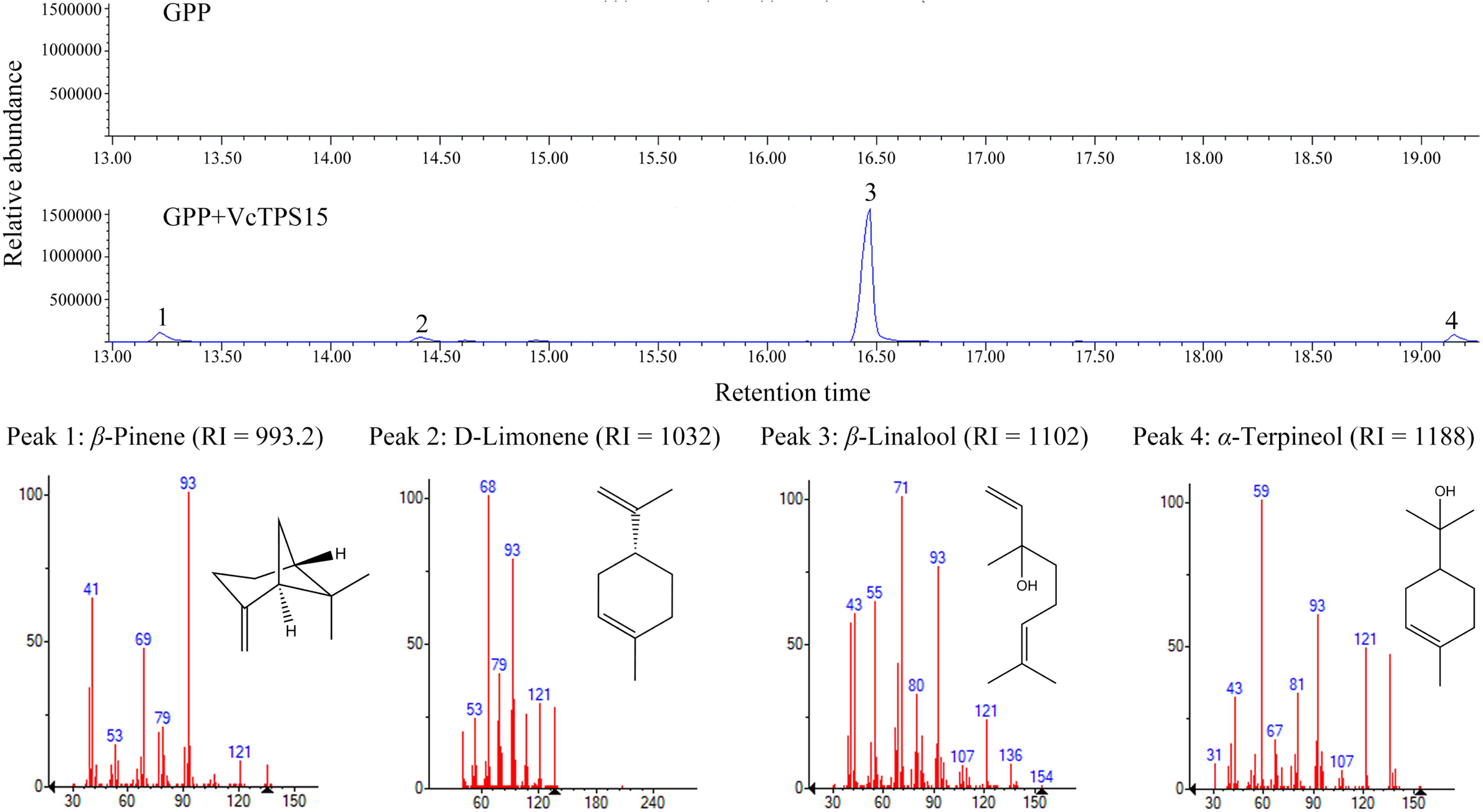
Selected ion chromatograms (m*/*z 93) of the products of the recombinant VcTPS15 protein using geranyl diphosphate (GPP) as substrate.

## 4. Conclusions

In this study, postharvest partial dehydration (∼30% weight loss) was applied to blueberries to improve the resulting blueberry wine sensory. Blueberry wines fermented from dehydrated blueberries had stronger sweet, floral and fruity aroma. The enhancement of sweet, floral and fruity aromas of blueberry wines fermented from dehydrated berries could be a consequence of the increases in phenylalanine-derived compounds such as phenylethanol, and terpenes such as linalool and geraniol, which were mainly derived from blueberries. The content of phenylalanine-derived compounds and terpenes in blueberries showed an increasing trend during postharvest dehydration process. Notably, postharvest dehydration upregulated the expressions of *VcGOT2* and *VcPAR*, leading to the significant increases in phenethyl alcohol level in blueberries. In addition, the upregulations of *VcDXS*, *VcHDR* and *VcTPS* could be linked to the higher terpene levels in dehydrated blueberries compared with controls. The function of the upregulated *VcTPS* was further characterized *in vitro*, it encoded a monoterpene synthase, catalyzing the production of linalool, β-pinene, D-limonene and α-terpineol from GPP. These finding provided insight into understanding the phenylalanine and terpene metabolism in blueberries in response to postharvest water loss, which laid theoretical basis for applying postharvest dehydration technology to improve blueberry wine sensory especially sweet, floral and fruity aromas. However, the underlying mechanisms regulating the key genes involved in phenylalanine and terpene metabolism in blueberries remains unknow, which should be further investigated in the future.

## Ethical approval

The sensory evaluation of blueberry wines in this study has been approved by the Ethics Committee of Anhui Agricultural University (Approval No.: KJLL2023020). Participants gave informed consent via the statement “I am aware that my responses are confidential, and I agree to participate in this sensory evaluation” where an affirmative reply was required to enter the sensory evaluation. They were able to withdraw from the sensory evaluation at any time without giving a reason.

## Acknowledgment

This work was supported by the Anhui Agricultural University Foundation for Stability and Introduction of Talent (rc352111), the Natural Science Research Project in Anhui Universities (2023AH051046), the Anhui Province Natural Science Foundation (Grant No. 2108085MC118), and the Special Fund for Anhui Province Academic and Technological Leader (2021D297).

## Author contributions

Yu Wang: Conceptualization; Supervision; Funding acquisition; Writing - Original draft. Qi Zhang, Investigation; Formal analysis; Visualization. Qin Yang, Investigation. Chen Bian, Investigation. Shu-Qin Huang, Investigation. Lu-Lu Zhao, Investigation. Ya-Qiong Huang, Investigation. Shan-Shan Shen, Formal analysis. Qi Chen, Formal analysis. Hai-Wei Zhang, Formal analysis. Xue-Ling Gao: Project administration; Funding acquisition; Writing - review & editing.

## Supplementary materials

**Table S1** Primers used for qRT-PCR.

**Table S2** Effects of postharvest dehydration on basic physiochemical parameters of blueberry, juice and wine.

**Table S3** Concentrations and relative odor activity values (rOAVs) of phenylalanine-derived compounds and terpene identified in blueberry wines fermented from fresh blueberries and partial dehydrated blueberries by HS-SPME-GC/MS.

**Table S4** Concentrations of phenylalanine-derived compounds and terpenes on a per fresh weight basis (μg/kg FW) in blueberries during postharvest dehydration process.

**Table S5** Concentrations of phenylalanine-derived compounds and terpenes on a per berry basis

(ng/berry) in blueberries during postharvest dehydration process.

**Table S6** The information of *VcTPS* genes identified in blueberries.

**Figure S1** Phylogenetic tree of full-length TPS from *Arabidopsis thaliana*, *Solanum lycopersicum*, *Vitis vinifera*, and *Vaccinium corymbosum*.

**Figure S2** Comparison of amino acid sequences of VcTPS15 in *Vaccinium corymbosum* and other TPS-g proteins in *Arabidopsis thaliana*, *Solanum lycopersicum* and *Vitis vinifera*.

**Figure S3** Expression profiles of *VcTPS* genes in different tissues during blueberry development. LD, leaf day. LN, leaf night. FL, flower at anthesis. PF, flower post-fertilization. Grnfrt, green blueberry fruit. Pinkfrt, pink blueberry fruit. Ripe, ripe blueberry fruit. Data sourced from http://gigadb.org/dataset/view/id/100537.

