## Supplementary material for "Postharvest partial dehydration of blueberries enhanced blueberry wine aroma via upregulating phenylalanine metabolism and terpene biosynthesis": Figure S1-S3

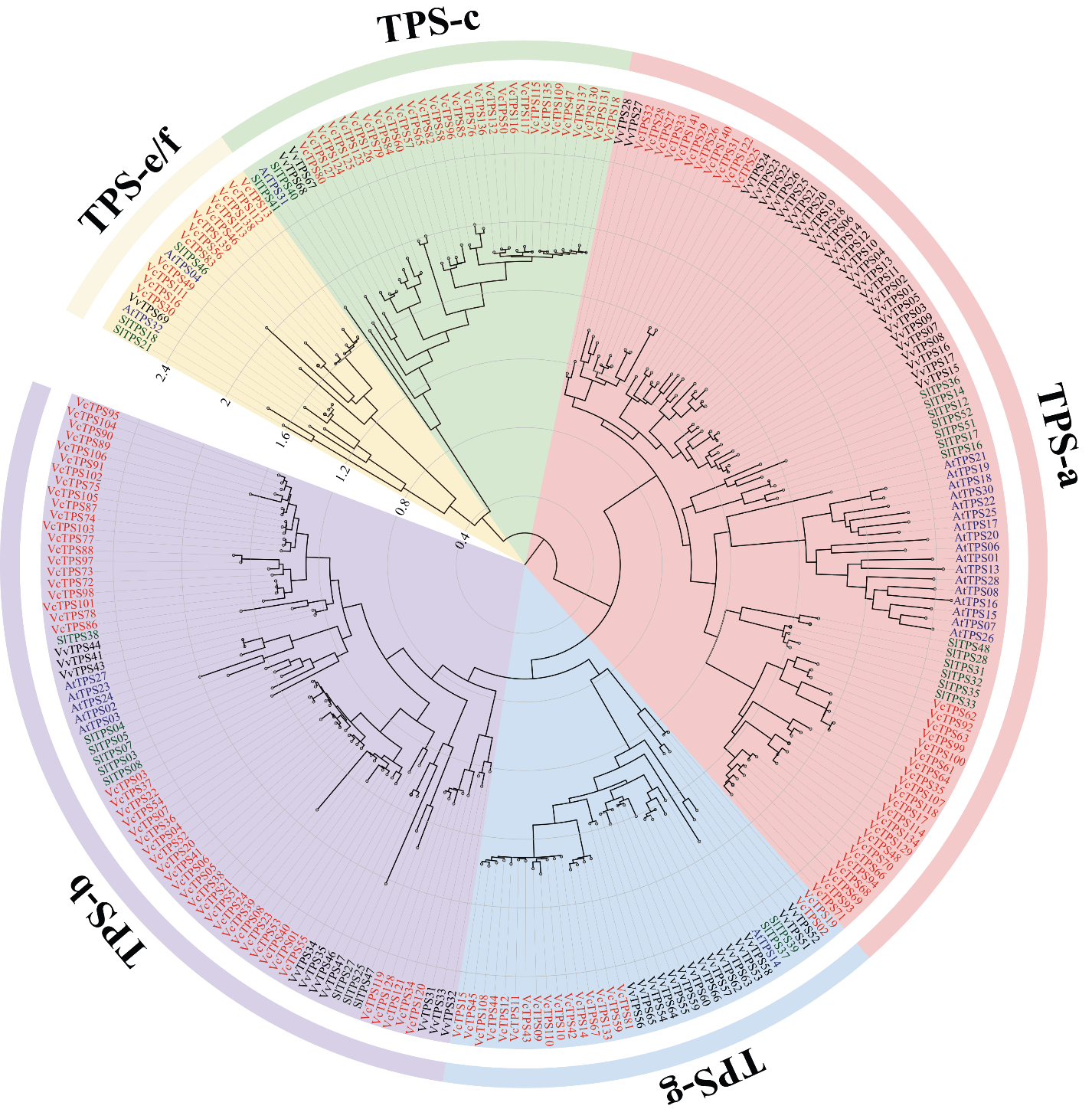
**Figure S1** Phylogenetic tree of full-length TPS from *Arabidopsis thaliana*, *Solanum lycopersicum*, *Vitis vinifera*, and *Vaccinium corymbosum*.

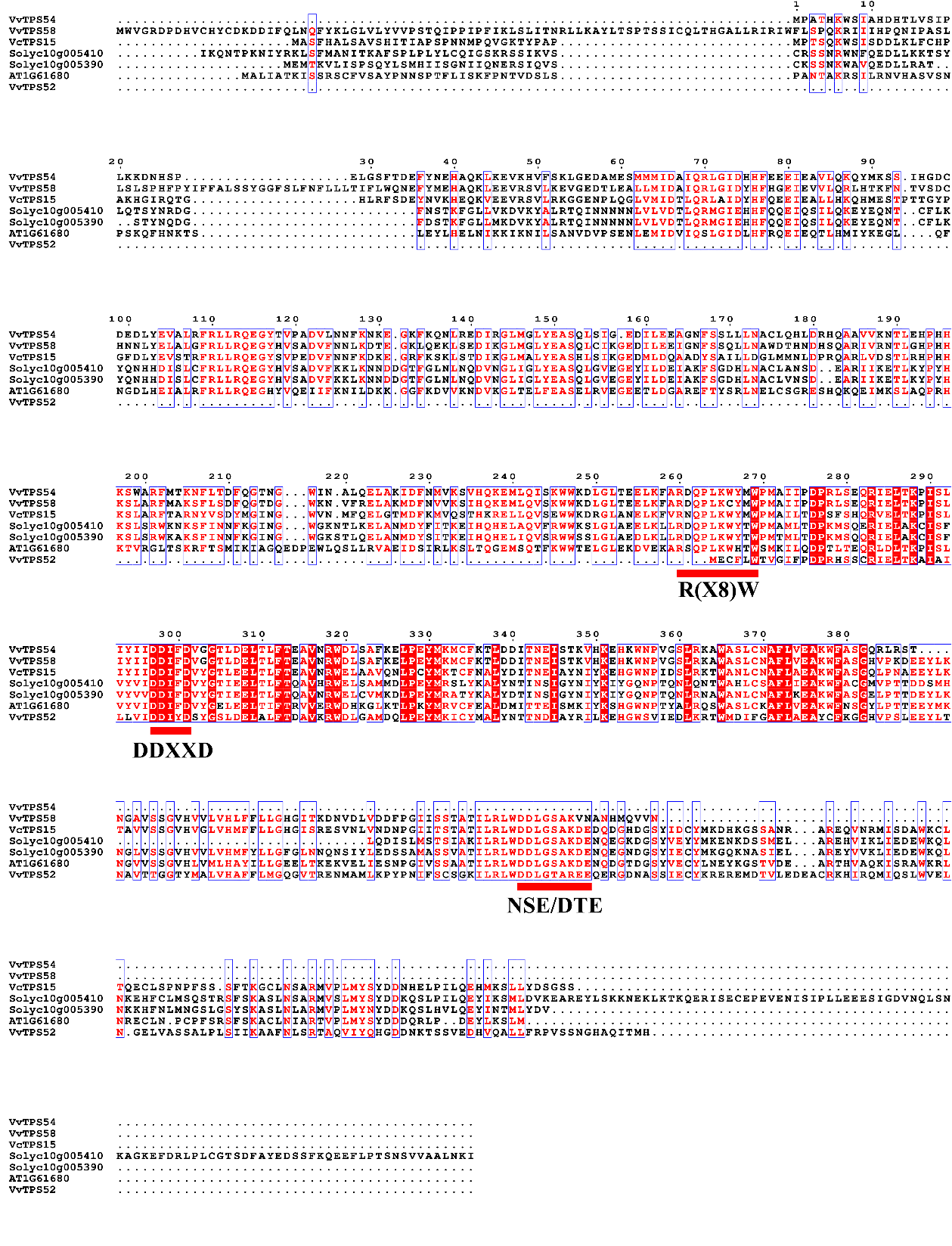
**Figure S2** Comparison of amino acid sequences of VcTPS15 in *Vaccinium corymbosum* and other TPS-g proteins in *Arabidopsis thaliana*, *Solanum lycopersicum* and *Vitis vinifera*.

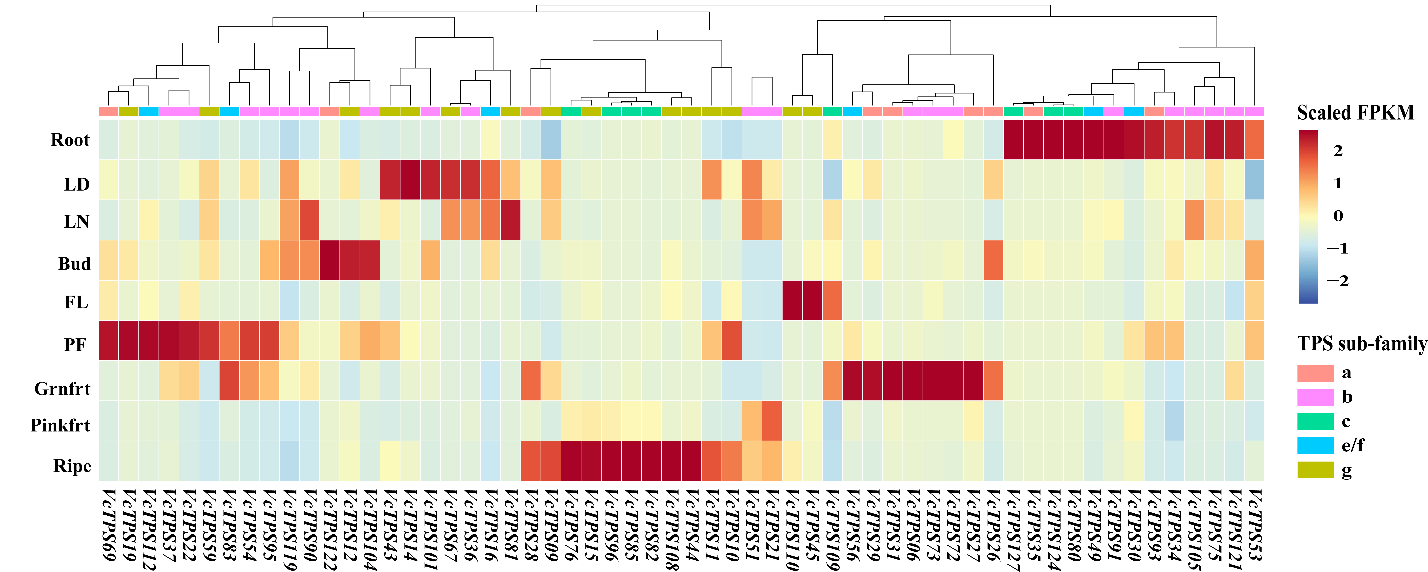
**Figure S3** Expression profiles of *VcTPS* genes in different tissues during blueberry development. LD, leaf day. LN, leaf night. FL, flower at anthesis. PF, flower post-fertilization. Grnfrt, green blueberry fruit. Pinkfrt, pink blueberry fruit. Ripe, ripe blueberry fruit. Data sourced from http://gigadb.org/dataset/view/id/100537.
