## Supplementary material for "Postharvest partial dehydration of blueberries enhanced blueberry wine aroma via upregulating phenylalanine metabolism and terpene biosynthesis": Table S1-S5

**Table S1** Primers used for RT-qPCR.

| **Gene** | **Gene ID** | **Forward primer (5’-3’)** | **Reverse primer (5’-3’)** |
| --- | --- | --- | --- |
| *VcDXS* | VaccDscaff1-snap-gene-85.24 | TGCTCCTCAGGACAGGCTAT | CTCAAAGCCCCTACTGGTGG |
| *VcHDR* | VaccDscaff2-augustus-gene-421.30 | TCGTGAAGAGCATGGCTGAG | GAATTCCACGATCCTCCGCA |
| *VcHDR* | VaccDscaff14-augustus-gene-366.23 | GCGATGTTGTGGTTTTGCCT | TTGTCGCTCTTGAGTGGCAT |
| *VcGGPS* | VaccDscaff12-augustus-gene-75.22 | GCCATGCGATACTCTCTGCT | ATCAGAAACTCCGGCGGAAG |
| *VcGGPS* | VaccDscaff23-snap-gene-336.32 | GCCTTCGCCTTCGAGTACAT | CACCTCGCGAACTTCCTCAA |
| *VcTPS* | VaccDscaff4-snap-gene-178.35 | TCTCGCAGCAAAGAGTGGAG | CAACTGCCCAGAAGCAAACC |
| *VcUBC28* | - | CCATCCACTTCCCTCCAGATTATCCAT | ACAGATTGAGAGCAGCACCTTGGA |

**Table S2** Effects of postharvest dehydration on basic physiochemical parameters of berry, juice and wine from fresh (CK) and partial dehydrated blueberries (WL30).

| **Sample type** | **Parameters** | **CK** | **WL30** |
| --- | --- | --- | --- |
| **Berry** | Total anthocyanins  (mg/kg FW) | 1727±40b ^b^ | 2153±80a |
|  | Total anthocyanins  (ng/berry) | 2210±52a | 1910±92b |
|  | Total phenols  (mg GAE/kg FW) | 3366±73b | 4567±198a |
|  | Total phenols  (ng GAE/berry) | 4308±93a | 4050±131a |
|  | Decay index ^a^ | 1 | 2 |
| **Juice** | Soluble solids(°Brix) | 14.57±0.15b | 18±0.17a |
|  | Sugar (g/L) | 159.2±4.9b | 196.3±3.5a |
|  | Titratable acidity (g/L) | 6.82±0.74a | 5.53±0.37a |
|  | pH | 3.10±0.01b | 3.23±0.05a |
|  | *L* | 42.88±1.2a | 44.79±2.14a |
|  | *a** | 56.39±0.88a | 53.27±1.47b |
|  | *b** | 19.68±2b | 24.18±0.43a |
| **Wine** | Residual sugar (g/L) | 4.76±0.57a | 4.48±0.41a |
|  | Titratable acidity (g/L) | 9.76±0.34a | 7.76±0.66b |
|  | pH | 3.34±0.05b | 3.44±0.02a |
|  | Total anthocyanins (mg/L) | 340.6±24.4b | 443.9±20.2a |
|  | Total phenols  (mg GAE/L) | 1833±47b | 2312±113a |
|  | Ethanol (%vol) | 11.75±0.99a | 12.69±0.23a |
|  | Total SO_2_ (mg/L) | 22.70±0.90a | 21.33±1.07a |
|  | *L* | 49.17±1.06b | 47.87±1.39a |
|  | *a** | 52.52±1.07b | 56.59±1.37a |
|  | *b** | 15.31±0.49b | 23.89±0.49a |

^a^ Decay index: no decay = 1; decay berries < 5% =2; decay berries < 10% = 3; decay berries < 20% = 4; decay berries > 20% = 5.

^b^ Different letters in the same row represent significant differences on the basis of Duncan’s multiple range test at *p* < 0.05.

**Table S3** Concentrations and relative odor activity values (rOAVs) of phenylalanine-derived compounds and terpene identified in blueberry wines fermented from fresh (CK) and partial dehydrated blueberries (WL30) by HS-SPME-GC/MS.

| **Compound** | **CAS** | **RI ^a^** | **Threshold ^b^  (μg/L)** | **Aroma descriptors** | **Concentrations (μg/L)** | |  | **rOAV** | |
| --- | --- | --- | --- | --- | --- | --- | --- | --- | --- |
|  |  |  |  |  | **CK** | **WL30** |  | **CK** | **WL30** |
| **Phenylalanine-derived compounds** | | | | | | |  |  |  |
| Methyl salicylate | 119368 | 1195.4 | 71 ^[1]^ | Peppermint | 5.22±1.96b ^c^ | 26.84±3.34a |  | < 0.1 | 0.38 |
| Benzaldehyde | 100527 | 960.5 | - | - | 0.81±0.18b | 51.26±6.02a |  | - | - |
| Phenylethanol | 60128 | 1112.2 | 14000 ^[2]^ | Rose, honey | 1280±106b | 1920±47a |  | < 0.1 | 0.14 |
| Phenethyl acetate | 103457 | 1255.6 | 250 ^[3]^ | Rose, honey | 45.27±2.91b | 66.58±2.67a |  | 0.18 | 0.27 |
| Ethyl phenylacetate | 101973 | 1240.6 | 40 ^[4]^ | Mint | 14.20±1.21b | 106.5±22.6a |  | 0.36 | 2.66 |
| Benzyl alcohol | 100516 | 987.7 | 200 ^[5]^ | Almonds | 4.34±0.72b | 6.68±0.93a |  | < 0.1 | < 0.1 |
| Phenylacetaldehyde | 122781 | 1021.6 | 1 ^[5]^ | Floral | 2.12±0.04b | 24.84±1.69a |  | 2.12 | 24.84 |
| **Terpenes** |  |  |  |  |  |  |  |  |  |
| Isoterpinolene | 586630 | 1013.1 | - | - | 0.10±0.01b | 1.29±0.08a |  | - | - |
| Sabinen | 3387415 | 988 | - | - | 26.03±0.84b | 31.82±2.17a |  | - | - |
| *α*-Terpinene | 99865 | 1013.6 | - | - | 1.80±0.30b | 13.80±1.33a |  | - | - |
| *o*-Cymene | 527844 | 1021.5 | - | - | 8.34±2.84b | 125.0±11.2a |  | - | - |
| D-Limonene | 5989275 | 1026 | - | - | 10.50±2.02a | 13.52±1.55a |  | - | - |
| *β*-Phellandrene | 555102 | 1025.7 | - | - | 10.74±2.22a | 13.54±1.73a |  | - | - |
| Eucalyptol | 470826 | 1026.4 | 1.1 ^[1]^ | Eucalyptus | 0.70±0.05b | 1.08±0.02a |  | 0.64 | 0.98 |
| *trans*-*β*-Ocimene | 3779611 | 1035.4 | 14 ^[2]^ | Apple peel, fruity | 4.85±0.49a | 5.20±0.97a |  | 0.35 | 0.37 |
| *β*-Ocimene | 13877913 | 1046 | - | - | 6.32±0.41a | 6.25±1.14a |  | - | - |
| *γ*-Terpinene | 99854 | 1055.8 | - | - | 1.25±0.10b | 8.35±1.89a |  | - | - |
| Acetophenone | 98862 | 1053.9 | - | - | Tr ^d^ | 1.72±0.39a |  | - | - |
| *cis*-Linalool oxide | 5989333 | 1068.7 | - | - | 1.68±0.05b | 4.00±0.45a |  | - | - |
| *allo*-Ocimene | 673847 | 1082.2 | - | - | 2.17±0.13b | 4.70±0.37a |  | - | - |
| Limonene epoxide | 1195922 | 1069 | - | - | 2.61±0.37a | 1.97±0.20a |  | - | - |
| Linalool | 78706 | 1099.8 | 15 ^[6]^ | Floral, lavender-like | 260.3±9.3b | 342.7±13.2a |  | 17.36 | 22.85 |
| Nerol oxide | 1786089 | 1149.8 | - | - | 0.55±0.02b | 0.89±0.09a |  | - | - |
| Rose oxide | 16409431 | 1108.4 | 0.2 ^[7]^ | Floral, rose | 2.99±0.41b | 5.75±0.56a |  | 14.95 | 28.75 |
| *endo*-Borneol | 507700 | 1168.6 | - | - | 3.76±0.39b | 12.96±0.77a |  | - | - |
| Terpinen-4-ol | 562743 | 1177.2 | 250 ^[8]^ | Turpentine, nutmeg | 6.31±3.50b | 403.9±27.1a |  | < 0.1 | 1.62 |
| *α*-Terpineol | 98555 | 1192.8 | 250 ^[9]^ | Lilac | 50.67±3.18b | 76.73±1.86a |  | 0.20 | 0.31 |
| *cis*-Geraniol | 106252 | 1222.8 | - | - | 11.14±2.43b | 18.51±1.45a |  | - | - |
| Citronellol | 106229 | 1226.1 | 100 ^[10]^ | Rose, citrus | 40.75±4.8b | 90.53±17.03a |  | 0.41 | 0.91 |
| Carvone | 99490 | 1241.5 | - | - | 49.96±9.93a | 20.38±3.17b |  | - | - |
| Geraniol | 106241 | 1249.5 | 36 ^[11]^ | Floral, citrus | 33.03±4.99b | 43.44±0.60a |  | 0.92 | 1.21 |
| *β*-Damascenone | 23726934 | 1443.8 | 0.007 ^[12]^ | Apple, rose, honey | 1.45±0.18b | 2.19±0.28a |  | 207.1 | 312.9 |
| Cedrol | 77532 | 1669.7 | - | - | 0.98±0.08a | 0.83±0.10a |  | - | - |

^a^ Retention indices on HP-5MS column.

^b^ Odor thresholds obtained from references.

^c^ Different letters in the same row means significant differences on the basis of Duncan’s multiple range test at *p* < 0.05.

^d^ Tr means trace concentration.

**Table S4** Concentrations of phenylalanine-derived compounds and terpenes on a per fresh weight basis (μg/kg FW) in blueberries during postharvest dehydration process.

| **Compound** | **CAS** | **RI ^a^** | **Concentrations (μg/kg FW)** | | | | | |
| --- | --- | --- | --- | --- | --- | --- | --- | --- |
|  |  |  | **0 d** | **2 d** | **4 d** | **6 d** | **8 d** | **10 d** |
| **Phenylalanine-derived volatiles** | | | | | | | | |
| Benzyl alcohol | 100516 | 988 | 3.42±0.36d ^b^ | 5.24±0.61cd | 8.29±0.94c | 11.77±0.43b | 14.07±1.83ab | 15.86±3.92a |
| Benzaldehyde | 100527 | 961 | 2.78±0.76c | 2.87±0.77c | 8.92±2.34c | 10.25±1.61c | 59.14±17.06b | 85.22±14.83a |
| Phenylethyl Alcohol | 60128 | 1112 | 12.11±2.28cd | 9.59±2.04d | 13.61±0.84cd | 22.25±7.21c | 37.84±0.67b | 48.79±10.89a |
| Ethyl benzeneacetate | 101973 | 1241 | Tr ^c^ | Tr | Tr | Tr | 10.14±2.68b | 16.95±2.83a |
| **Terpenes** | | | | | | | | |
| *β*-Pinene | 127913 | 986 | 79.8±17.5d | 91.7±8.4cd | 101.9±2.2bcd | 112.4±8.8bc | 125.2±10.8b | 155.9±31.6a |
| *α*-Phellandrene | 4221981 | 1012 | 10.25±2.63c | 12.47±1.48c | 14.63±1.89c | 21.32±4.53b | 23.12±4.61ab | 27.72±3.41a |
| *α*-Terpinene | 99865 | 1014 | 6.3±1.43d | 11.23±2.7c | 19.4±0.84b | 23.06±3.64ab | 26.18±0.7a | 26.55±2.66a |
| Terpineol | 8000417 | 1019 | 1.72±0.43c | 1.97±0.47c | 2.42±0.93bc | 2.87±0.31bc | 3.33±0.97b | 4.65±0.57a |
| *o*-Cymene | 527844 | 1022 | 52.85±3.15c | 64.48±1.54c | 180.3±15.5b | 216.9±25.6ab | 231.9±25.8a | 235.4±32.4a |
| D-Limonene | 5989275 | 1026 | 20.41±0.97e | 22.66±0.38de | 24.73±1.16cd | 26.58±0.42c | 31.46±1.72b | 57.93±4.48a |
| *trans*-*β*-Ocimene | 3779611 | 1035 | 6.94±0.91e | 13.9±1.03d | 15.47±1.86cd | 18.1±1.13bc | 19.07±0.83b | 24.23±2.53a |
| *β*-Ocimene | 13877913 | 1046 | 16.85±1.86c | 17.42±2.84c | 23.7±1.12b | 25.03±1.79b | 25.02±2.59b | 30.69±1.8a |
| *trans*-Linalool oxide (furanoid) | 34995772 | 1068 | 2.94±0.47b | 2.87±0.69b | 3.17±0.45b | 3.31±0.17b | 4.1±0.26a | 4.82±0.25a |
| Linalool | 78706 | 1100 | 44.71±3.5f | 55.71±3.81e | 67.25±5.37d | 76.97±1.73c | 91.23±4.64b | 131.5±4.8a |
| Camphor | 76222 | 1143 | 2.72±0.14a | 2.05±0.18b | 1.68±0.14c | 1.73±0.18c | 1.73±0.24c | 1.36±0.04d |
| Pinocarveol | 5947364 | 1124 | 1.86±0.26c | 2.42±0.66bc | 2.02±0.31c | 2.52±0.22bc | 3.15±0.8ab | 3.78±0.76a |
| *endo*-Borneol | 507700 | 1169 | 142.0±4.6d | 164.6±3.5c | 171.7±17.2bc | 173.1±4.9bc | 185.8±7.7b | 218.6±6.5a |
| Menthol | 89781 | 1172 | 2.58±0.3c | 4.04±1.1b | 4.12±0.84b | 4.71±0.89ab | 4.52±0.35ab | 5.65±0.41a |
| Terpinen-4-ol | 562743 | 1177 | 19.22±1.97abc | 16.4±1.18cd | 15.99±0.9d | 17.55±1.39bcd | 20.55±2.07a | 20.32±1.28ab |
| *α*-Terpineol | 98555 | 1193 | 74±4.51c | 83.17±2.19bc | 93.39±2.98b | 114.5±17.4a | 95.3±1.78b | 125.7±15.3a |
| *cis*-*β*-Farnesene | 28973979 | 1450 | 1.82±0.53d | 2.99±0.44c | 4.53±0.71b | 4.77±0.71b | 5.01±0.96b | 7.27±0.41a |
| *β*-Damascenone | 23726934 | 1444 | 155.7±8.2d | 302.2±29.7a | 316.1±13.5a | 232.0±21.9c | 270.5±5.9b | 244.0±2.2bc |
| *cis*-Geraniol | 106252 | 1223 | 9.97±2.97d | 18.62±0.02c | 19.81±3.5bc | 18.86±2.62c | 25.15±5.14b | 33.27±1.86a |
| Geraniol | 106241 | 1250 | 109.2±35.7d | 270.2±35.6b | 211.6±12.2c | 246.2±19.3bc | 253.5±4.5bc | 330.6±6.4a |

^a^ Retention indices on HP-5MS column.

^b^ Different letters in the same row means significant differences on the basis of Duncan’s multiple range test at *p* < 0.05.

^c^ Tr means trace concentration.

**Table S5** Concentrations of phenylalanine-derived compounds and terpenes on a per berry basis (ng/berry) in blueberries during postharvest dehydration process.

| **Compound** | **CAS** | **RI** | **Concentrations (ng/berry)** | | | | | |
| --- | --- | --- | --- | --- | --- | --- | --- | --- |
|  |  |  | **0 d** | **2 d** | **4 d** | **6 d** | **8 d** | **10 d** |
| **Phenylalanine-derived compounds** | | | | | | | | |
| Benzyl alcohol | 100516 | 988 | 4.38±0.46c | 6.51±0.77c | 9.75±1.1b | 12.95±0.43a | 13.99±1.79a | 14.07±3.45a |
| Benzaldehyde | 100527 | 961 | 3.56±0.97c | 3.57±0.97c | 10.5±2.75c | 11.27±1.77c | 58.87±17.25b | 75.6±13.26a |
| Phenylethyl alcohol | 60128 | 1112 | 15.5±2.92bc | 11.91±2.53c | 16.01±0.98bc | 24.49±8b | 37.64±0.49a | 43.22±9.19a |
| Ethyl benzeneacetate | 101973 | 1241 | Tr ^c^ | Tr | Tr | Tr | 10.09±2.7b | 15.02±2.36a |
| **Terpenes** | | | | | | | | |
| *β*-Pinene | 127913 | 986 | 102.2±22.4b | 113.9±10.7ab | 119.8±2.5ab | 123.6±10.1ab | 124.5±11.3ab | 138.1±26.2a |
| *α*-Phellandrene | 4221981 | 1012 | 13.12±3.37c | 15.48±1.83c | 17.21±2.22bc | 23.45±5.04ab | 23±4.58ab | 24.57±2.72a |
| *α*-Terpinene | 99865 | 1014 | 8.06±1.84c | 13.94±3.33b | 22.82±1a | 25.37±4.03a | 26.04±0.8a | 23.55±2.33a |
| Terpineol | 8000417 | 1019 | 2.2±0.55b | 2.44±0.58b | 2.85±1.1ab | 3.16±0.34ab | 3.31±0.95ab | 4.12±0.47a |
| *o*-Cymene | 527844 | 1022 | 67.7±4.0b | 80.1±1.9b | 212.1±18.3a | 238.5±28.1a | 230.7±26.3a | 208.8±28.6a |
| D-Limonene | 5989275 | 1026 | 26.12±1.25c | 28.13±0.5bc | 29.1±1.36bc | 29.23±0.39bc | 31.28±1.58b | 51.37±3.45a |
| *trans*-*β*-Ocimene | 3779611 | 1035 | 8.89±1.16c | 17.25±1.26b | 18.2±2.19b | 19.9±1.17ab | 18.97±0.74ab | 21.5±2.42a |
| *β*-Ocimene | 13877913 | 1046 | 21.57±2.38b | 21.63±3.51b | 27.88±1.31a | 27.52±1.92a | 24.88±2.53ab | 27.22±1.39a |
| *trans*-Linalool oxide (furanoid) | 34995772 | 1068 | 3.77±0.6a | 3.57±0.85a | 3.73±0.53a | 3.64±0.19a | 4.08±0.25a | 4.27±0.2a |
| Linalool | 78706 | 1100 | 57.22±4.48e | 69.18±4.87d | 79.11±6.28c | 84.65±2.01bc | 90.75±5.04b | 116.7±4.7a |
| Camphor | 76222 | 1143 | 3.48±0.17a | 2.54±0.22b | 1.98±0.16c | 1.91±0.2c | 1.72±0.23c | 1.21±0.05d |
| Pinocarveol | 5947364 | 1124 | 2.38±0.33a | 3±0.81a | 2.38±0.37a | 2.77±0.25a | 3.13±0.78a | 3.36±0.69a |
| *endo*-Borneol | 507700 | 1169 | 181.7±5.8c | 204.4±4.6a | 202.0±20.3ab | 190.3±4.9abc | 184.7±6.7bc | 193.9±4.0abc |
| Menthol | 89781 | 1172 | 3.31±0.38b | 5.01±1.36a | 4.85±0.99ab | 5.18±0.98a | 4.49±0.32ab | 5.02±0.41a |
| Terpinen-4-ol | 562743 | 1177 | 24.6±2.53a | 20.37±1.51b | 18.81±1.06b | 19.3±1.58b | 20.44±2.02b | 18.02±0.95b |
| *α*-Terpineol | 98555 | 1193 | 94.72±5.78b | 103.3±2.9b | 109.9±3.5ab | 125.9±18.7a | 94.8±2.2b | 111.5±12.4ab |
| *cis*-*β*-Farnesene | 28973979 | 1450 | 2.33±0.67d | 3.71±0.55c | 5.33±0.84ab | 5.24±0.78ab | 4.99±0.95bc | 6.45±0.33a |
| *β*-Damascenone | 23726934 | 1444 | 199.3±10.5c | 375.3±37.6a | 371.9±16.0a | 255.1±23.6b | 269.0±6.6b | 216.5±4.3c |
| *cis*-Geraniol | 106252 | 1223 | 12.76±3.8c | 23.11±0.07ab | 23.31±4.11ab | 20.73±2.81b | 25.02±5.16ab | 29.53±1.9a |
| Geraniol | 106241 | 1250 | 139.8±45.8c | 335.5±43.9a | 249.0±14.5b | 270.7±20.5b | 252.1±3.3b | 293.3±6.8ab |

^a^ Retention indices on HP-5MS column.

^b^ Different letters in the same row means significant differences on the basis of Duncan’s multiple range test at *p* < 0.05.

^c^ Tr means trace concentration.
